# Transcriptional analysis of efferocytosis in mouse skin wounds

**DOI:** 10.1101/2024.08.12.607219

**Authors:** Will Krause, Diane King, Valerie Horsley

**Affiliations:** Dept. of Molecular, Cellular, and Developmental Biology, Yale University, New Haven, Connecticut, USA; SunnyCrest Bioinformatics, Flemington, New Jersey, USA; Dept. of Dermatology, Yale School of Medicine, New Haven, Connecticut, USA

## Abstract

Defects in apoptotic cell clearance, or efferocytosis, can cause inflammatory diseases and prevent tissue repair due in part to inducing a pro-repair transcriptional program in phagocytic cells like macrophages. While the cellular machinery and metabolic pathways involved in efferocytosis have been characterized, the precise efferocytic response of macrophages is dependent on the identity and macromolecular cues of apoptotic cells, and the complex tissue microenvironment in which efferocytosis occurs. Here, we find that macrophages undergoing active efferocytosis in mid-stage mouse skin wounds in vivo display a pro-repair gene program, while efferocytosis of apoptotic skin fibroblasts in vitro also induces an inflammatory transcription response. These data provide a resource for understanding how the skin wound environment influences macrophage efferocytosis and will be useful for future investigations that define the role of efferocytosis during tissue repair.

## Introduction

The clearance of apoptotic cells, called efferocytosis, is required for tissue development, homeostasis, and repair to prevent harmful release of cellular debris from apoptotic cells (Freire-de-Lima et al., 2006; Waterborg et al., 2018; Zhang et al., 2019). Defects in efferocytosis are associated with autoimmune and inflammatory diseases which prolong inflammation and tissue damage, leading to organ dysfunction in several diseases including systemic autoimmune disorders, neurodegenerative, lung, gastrointestinal and heart diseases, and defects in wound healing (Boada-Romero et al., 2020). Inhibition of efferocytosis machinery can prevent tissue repair in mice during atherosclerosis (Kiss et al., 2006; Wang et al., 2017; Yurdagul et al., 2020), myocardial infarction (Bossi et al., 2014; Zhang et al., 2019) rheumatoid arthritis (Waterborg et al., 2018) and skin wound healing (Justynski et al., 2023). Interestingly, efferocytosis defects can also promote anti-tumor immunity in mice (Boada-Romero et al., 2020). Thus, understanding how efferocytosis controls inflammation has broad implications for tissue biology and disease prevention.

The primary effectors of efferocytosis are phagocytic immune cells such as macrophages and dendritic cells (Boada-Romero et al., 2020). After injury, infiltrating inflammatory macrophages perform efferocytosis, which stimulates signaling and metabolic changes that suppress inflammation and promote a resolving, tissue repair phenotype. Efferocytosis of apoptotic cells by phagocytic macrophages in vitro promotes the expression of anti-inflammatory cytokines (Fadok et al., 1998) as well as enzymes that produce pro-resolving lipid mediators (Dalli & Serhan, 2012), and in concert with IL-4 stimulation, a tissue repair gene signature (Bosurgi et al., 2017). Beyond changes in macrophage gene expression, apoptotic cell-derived DNA can enhance resolution of inflammation by promoting basal and resolving macrophage proliferation (Gerlach et al., 2021). Interestingly, the cellular source of apoptotic cells can drive macrophage heterogeneity to distinct gene expression programs and myeloid functions in a murine hepatic infection model (Liebold et al., 2024). Tissue specific cues can also regulate macrophage phenotype associated with efferocytosis (Gonzalez et al., 2017). Thus, the mechanisms by which efferocytosis impacts inflammation are still not fully defined and are likely dependent on several factors within the complex environments of specific tissues including the apoptotic cell identity and composition.

Here, we use a genetic mouse model to define how efferocytosis of non-myeloid cells impacts macrophage transcription within skin wounds. We find that efferocytic macrophages in mouse skin wounds upregulate pathways involved in angiogenesis and inflammatory processes, which are distinct from efferocytosis pathways induced by apoptotic fibroblasts in vitro. Finally, we examine how efferocytosis gene signatures define macrophage heterogeneity within single cell RNA sequencing data from skin wounds. These data reveal a distinct resolving transcriptional program associated with macrophage efferocytosis within mid-stage mouse skin wounds and will be useful for future investigations that define the role of efferocytosis during tissue repair.

## Results

### Isolation and genomic analysis of efferocytosis in skin wounds using LysMCreER/mTmG mice

To analyze macrophage efferocytosis during skin wound repair in vivo, we sought to determine whether genetic labeling of macrophages using LysMCreER mice crossed to reporter mice expressing mTmG could detect macrophage-driven efferocytosis in vivo (**Fig. 1a**). In LysMCreER/mTmG mice, intraperitoneal injection of tamoxifen induces a switch from membrane TdTomato (TdTom) expression to membrane GFP expression (mGFP) in *Lysozyme 2* (*LysM*) expressing myeloid cells, including macrophages and dendritic cells (Muzumdar et al., 2007; Shook et al., 2018), (**Fig. 1b**). Monocyte-derived macrophages are a prevalent phagocytic cell within early skin wounds (Aitcheson et al., 2021; Shook et al., 2018; Willenborg et al., 2012) and thus we analyzed mGFP expression in skin wounds of tamoxifen-treated LysMCreER/mTmG mice at D1 and D3 after injury. D1 wounds contained just ∼20% F4/80+CD45+ cells while D3 wounds contained ∼80% F4/80+CD45+ cells, suggesting D1 wounds had fewer mature macrophages (**Fig. S1, Fig. S2a**). D1 wounds contained ∼30% CD45+ F4/80+ macrophages that were GFP+ and this percentage increased to 85.8% in D3 skin wounds (**Fig. 1c, 1d, and S1, S2b**). The percentage of CD45+ F4/80− cells that were GFP+ decreased from 60% to 30% from D1 to D3, which likely reflects the loss of labelled neutrophils (Abram et al., 2014) at these wound stages (Wasko et al., 2022) (**Fig. 1c, 1d, Fig.S1, and S2b**).

**Figure 1:**
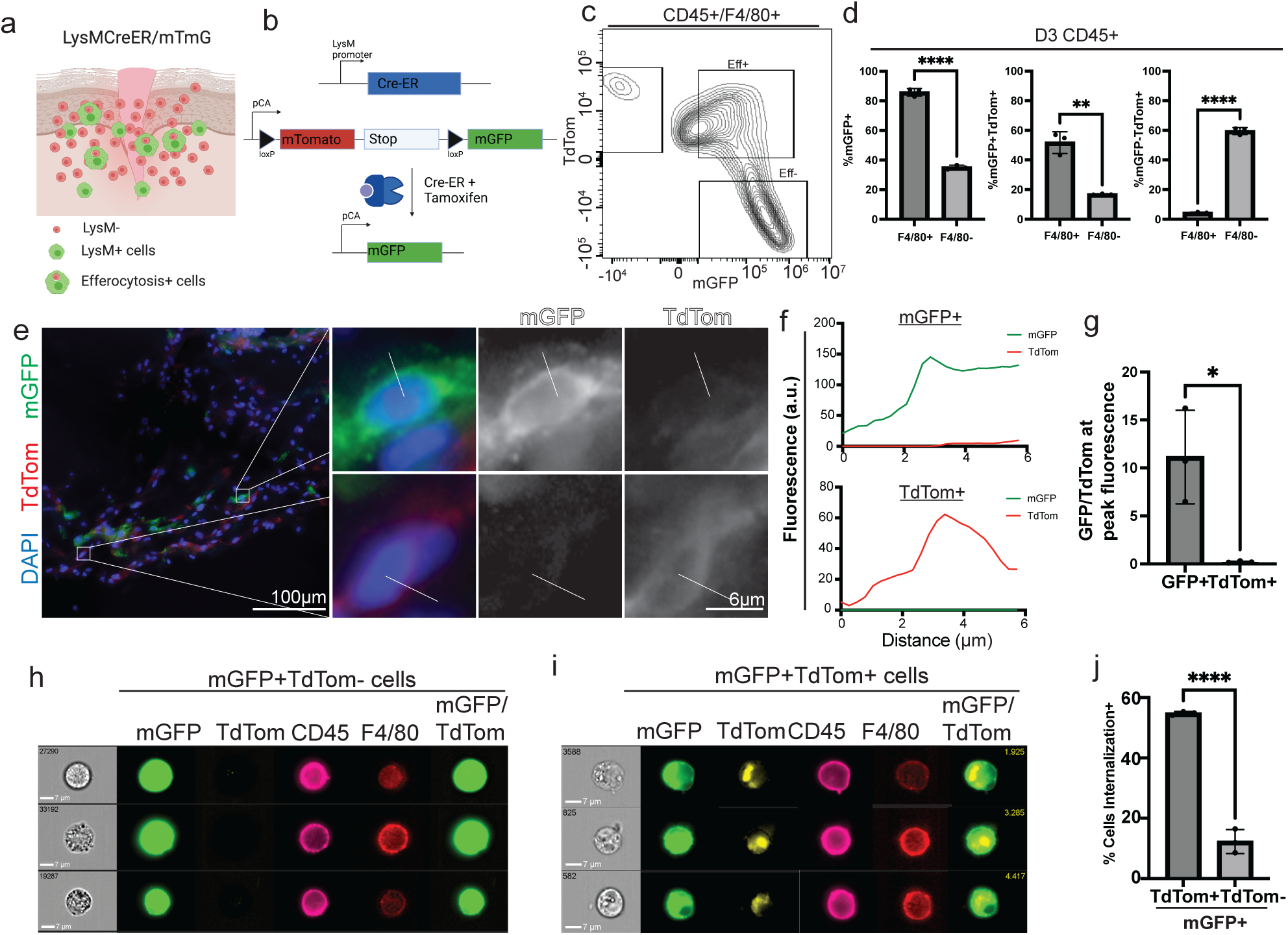
LysMCreER/mTmG mouse wounds contain efferocytosis+ macrophages. a-b) Schematic of *LysMCreER*/mTmG mouse wounds and genetic scheme. c) Density plots of AMNIS flow cytometry of F4/80+CD45+ macrophages (Macs) with mGFP and TdTom expression from day 3 wounds of *LysMCreER*/mTmG mice. d) Quantification of flow cytometry data for %GFP+, %GFP+TdTom+, and %GFP+TdTom− of F4/80+/CD45+ macrophages and F4/80−/CD45+ immune cells. n= average for 3 mice. Error bars indicate mean +/− SEM. p values calculated with one way ANOVA. e) Representative images from day 3 wounds of LysMCreER/mTmG mice. Line indicates line scan for quantification in g. f) Representative quantification of line scans (as shown in e) of fluorescence in GFP+ and TdTom+ cells in day 3 wounds of LysMCreER/mTmG mice. g) Quantification of the ratio of GFP/TdTom fluorescence at fluorescent peak for GFP+ or TdTom+ cells in day 3 wounds of LysMCreER/mTmG mice. At least 25 different cells were quantified for each type. N= 3 mice. Error bars indicate mean +/− SEM. p values calculated with one way ANOVA. h) Representative images of GFP+/TdTom− cells from AMNIS flow cytometry. i) Representative images of GFP+/TdTom+ cells from AMNIS flow cytometry. j) Quantification of % the GFP+ cells with internalized TdTom. n= 3 mice with at least 4500 cells analyzed. Error bars indicate mean +/− SEM. p values calculated with one way ANOVA.

We reasoned that GFP+ cells performing efferocytosis of *LysM* negative cells would also contain TdTom+ fluorescence (**Fig. 1a**). Analysis of TdTom in GFP+ cells of D1 and D3 wounds of LysMCreER/mTmG mice identified ∼20% and ∼45% of CD45+ F4/80+ macrophages that were GFP+TdTom+, respectively (**Fig. 1d, Fig. S2c**). In D1 wounds, about 35% of CD45+ F4/80+ cells were GFP− and TdTom+, and this percentage decreased to 4.25% by D3 (**Fig. 1d, Fig. S2d**). These data indicated that D3 wound beds of tamoxifen-treated LysMCreER/mTmG mice display efficient *LysM* Cre activity in our induction paradigm, and some GFP+ macrophages harbor TdTom expression.

Double positive mGFP and TdTom macrophages could be either mGFP+ cells undergoing efferocytosis or mGFP+ cells that retain TdTom expression. To distinguish between these possibilities, we used fluorescence microscopy to analyze mGFP+ and TdTom+ cells in sections of D3 skin wounds from LysMCreER/mTmG mice. To analyze mGFP and TdTom fluorescence in cell membranes of individual cells, we measured the fluorescence of GFP and TdTom in GFP+ and TdTom+ cells using line scans from the middle of the cell nucleus to the cell periphery (**Fig. 1e**). We found that GFP+ cells displayed a peak of GFP fluorescence outside of the nucleus with low TdTom fluorescence throughout the cell. TdTom+ cells displayed a similar trend with a peak of TdTom expression with low levels of GFP expression (**Fig. 1f**). Quantification of the ratio of peak GFP fluorescence in wounds of multiple mice confirmed that this trend was consistent (**Fig. 1g**).

For a more quantitative view of fluorescence localization, we utilized AMNIS flow cytometry, which combines conventional flow cytometry with microscopy to produce fluorescent images of each cell (Basiji, 2016). Cells from D3 wounds of tamoxifen-treated LysMCreER/mTmG mice were imaged for GFP, TdTom, CD45, and F4/80 expression. We found that mGFP+ TdTom− cells expressed F4/80, and that in mGFP+ cells, TdTom was localized within the cell rather than on the cell surface like CD45 and F4/80. To quantify the internalization of TdTom, we created a cell membrane mask and quantified internalization of the TdTom fluorescent signal. We found that 54.7% of CD45+ F4/80+ mGFP+ TdTom+ cells contained internalized TdTom signal (**Fig. 1h-j**), which is consistent with the quantification of our FACS data (**Fig. 1d**).

Having confirmed that macrophages from wounds of LysMCreER/mTmG mice contained internalized TdTom, we sought to use this system to analyze how efferocytosis impacts macrophage gene expression in skin wounds in vivo. We FACS purified live, F4/80+, GFP+ macrophages that were either TdTom+ (Eff+) or TdTom− (Eff−) from D3 wounds of 3 tamoxifen-treated LysMCreER/mTmG mice (**Figs. 2a, 2b, and S3**).

**Figure 2:**
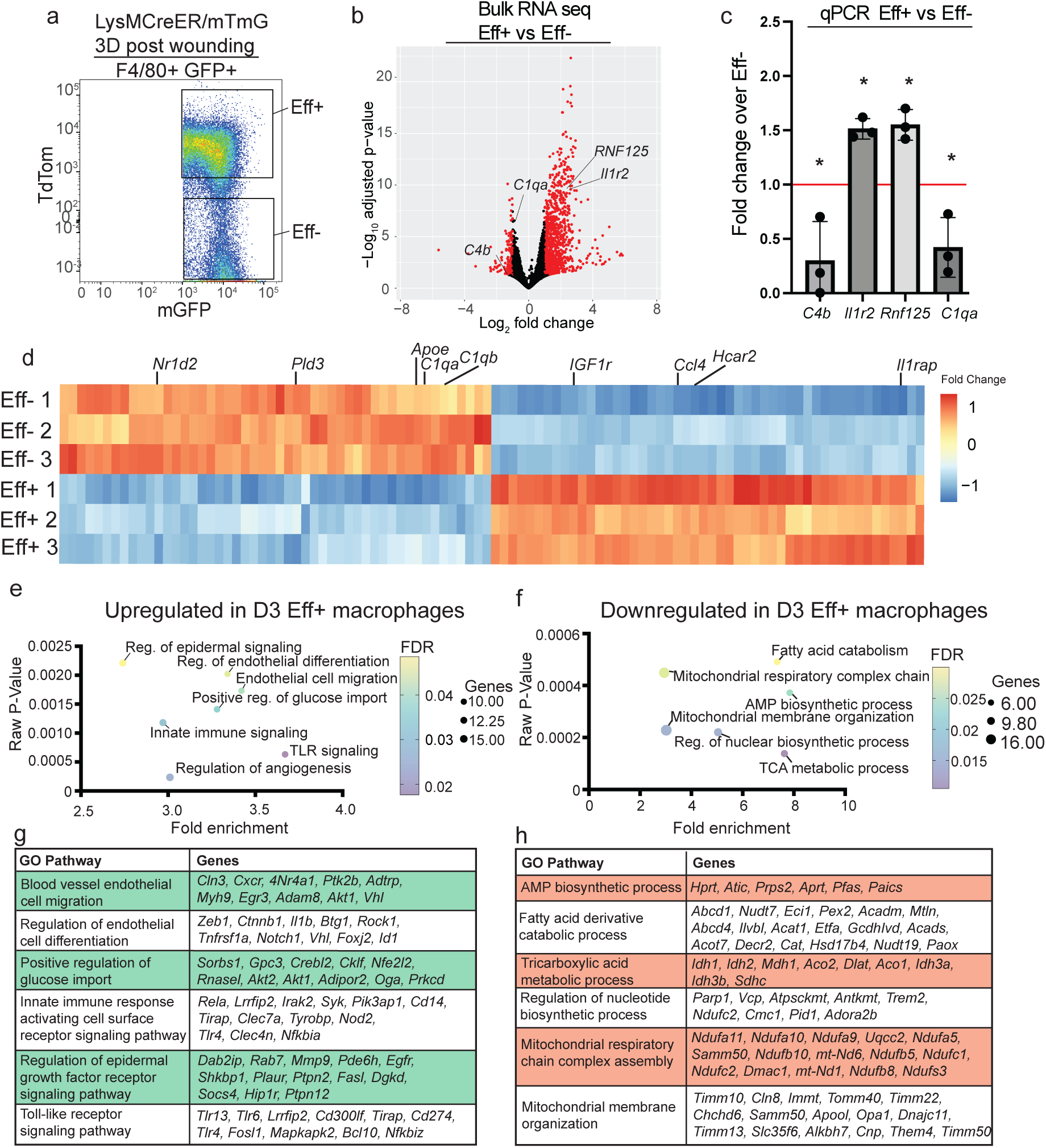
Transcriptional response of efferocytosis+ macrophages in mid-stage mouse skin wounds. a) Dot plot used efferocytosis+ (Eff+) and efferocytosis− (Eff−) F4/80+ macrophages. b) Volcano plot of differentially regulated genes Eff+ and Eff− macrophages. c) qPCR of mRNAs upregulated in sorted Eff+ macrophages compared to Eff− macrophages. N=3 mice. Error bars indicate mean +/− SEM. p values calculated with one way ANOVA. d) Heatmap of differentially regulated genes, showing top 50 upregulated and top 50 downregulated genes in Eff+ cells compared to Eff− cells by adjusted p-value. e-f) Bubble plot showing selected gene ontology terms from PantherGO analysis of genes upregulated (e) or downregulated (f) in Eff+ compared to Eff−. g-h) List of genes shown in the GO terms in (e) and (f).

Comparted to Eff− macrophages, Eff+ macrophages upregulated 1897 genes significantly, and downregulated 1147 genes significantly (**Fig. 2b and Table S1**). Selected upregulated and downregulated genes were validated by qPCR (**Fig. 2c**). The top 100 differentially expressed genes were consistently different between the 3 biological replicates (**Fig. 2d**). Gene pathway enrichment analysis of upregulated and downregulated genes in differentially expressed in Eff+ macrophages compared to Eff− macrophages was completed using Gene Ontology (GO) biological processes. This analysis revealed that Eff+ macrophages upregulated genes associated with angiogenesis, innate immune signaling, and epidermal growth factor signaling (**Figs. 2e and 2g, Table S3**). By contrast, Eff+ macrophages downregulated genes associated with the adenosine monophosphate (AMP) biosynthetic pathway, metabolic, and the nucleotide biosynthetic pathways (**Fig. 2f, 2h, Table S4**).

### Analysis of apoptotic fibroblast efferocytosis in vitro

During skin wound healing, different cell types undergo apoptosis and other forms of cell death, and caspase genes are expressed in multiple cell types including but not limited to dendritic cells, neutrophils, and fibroblasts (Justynski et al., 2023). In particular, we were intrigued that fibroblasts from early wounds expressed apoptotic markers (Justynski et al., 2023). Thus, we sought to determine whether macrophage efferocytosis of fibroblasts induced a specific efferocytosis signature as has been shown for other cell types (Liebold et al., 2024). To this end, we induced TdTom+ primary mouse fibroblasts from mTmG mice to undergo apoptosis by UV irradiation and incubated them with macrophages from WT mice that were differentiated from WT bone marrow derived monocytes (BMDMs) (**Fig. 3a**). After 12 hours, we FACS purified F4/80+ TdTomato+ macrophages (Eff+) and F4/80+ TdTomato− macrophages (Eff−) (**Fig. S4a**). We found that 80% of F4/80+ cells were TdTom− and only 20% were TdTom+ (**Fig. S4b**).

**Figure 3:**
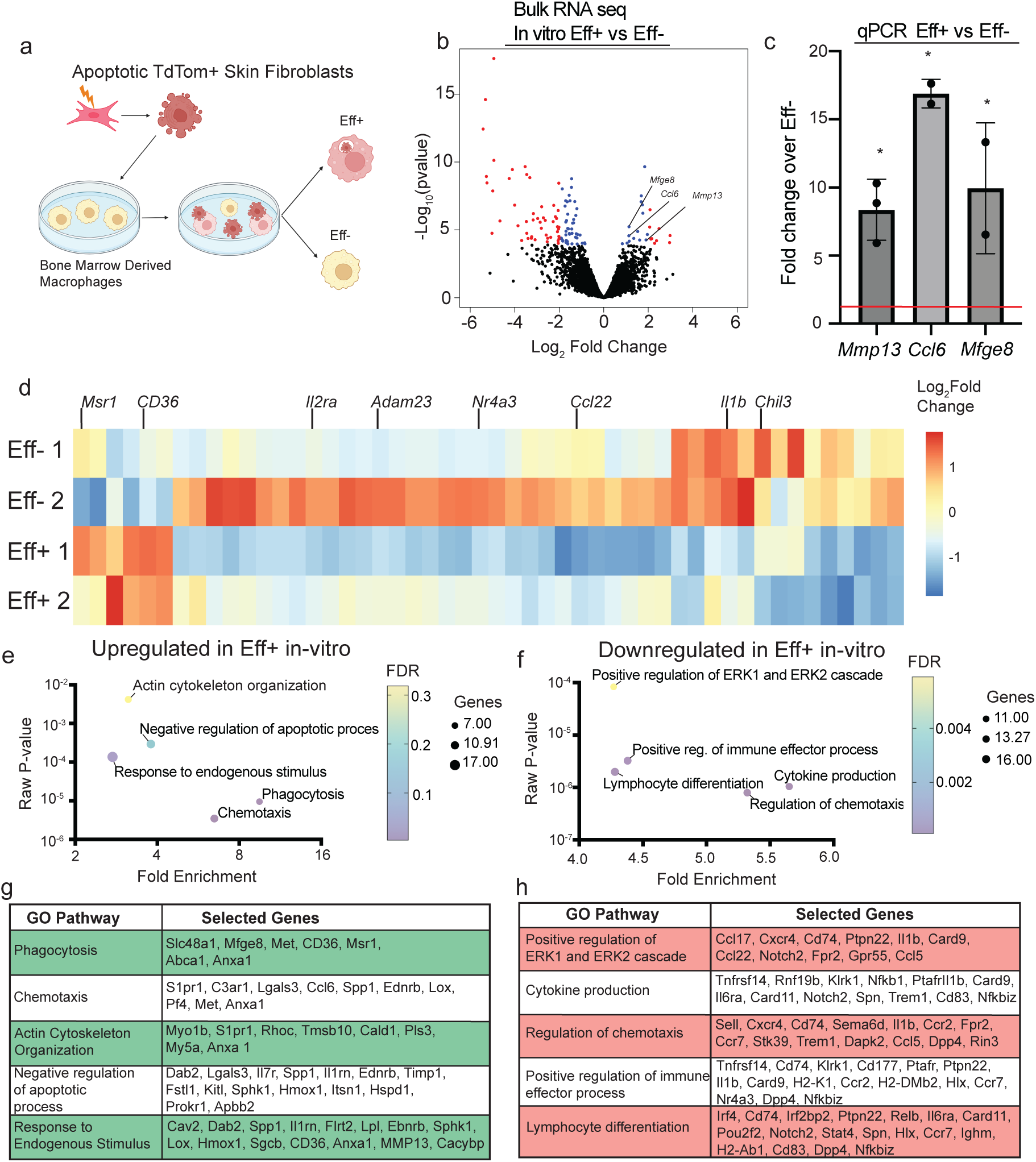
Transcriptional response of macrophage efferocytosis of apoptotic fibroblasts in vitro. a) Schematic of in vitro experiment. b) Volcano plot of differentially regulated genes. N=2 repeat experiments. c) qPCR for selected genes upregulated in Eff+ cells compared to Eff−cells treated with apoptotic fibroblasts. N=3 experiments. Error bars indicate mean +/− SEM. p values calculated with one way ANOVA. d) Heatmap of the top 50 differentially regulated genes in Eff+ cells compared to Eff− cells treated with apoptotic fibroblasts by adjusted P-value. e-f) Bubble plots showing selected gene ontology terms from differentially regulated genes in Eff+ cells compared to Eff− cells treated with apoptotic fibroblasts. g-h) Gene lists from selected pathways in e and f.

To evaluate gene expression changes between Eff+ macrophages and Eff− macrophages that were incubated with apoptotic fibroblasts, we performed bulk RNA sequencing. 103 genes were significantly upregulated, and 267 genes were significantly downregulated in Eff+ macrophages compared to Eff− macrophages (**Fig. 3b and Table S2**). Selected genes were validated by qPCR (**Fig. 3c**) and the top 50 differentially regulated genes were consistent across replicates (**Fig. 3d**). Gene ontology analysis of differentially regulated genes revealed that typical efferocytosis pathways: phagocytosis, chemotaxis, and actin cytoskeleton organization pathways were upregulated in Eff+ **(Figs. 3e and 3g, Table S5**), while inflammatory cytokine, lymphocyte differentiation, and metabolic pathways were downregulated (**Fig. 3f, 3h, and Table S6**).

### Analysis of efferocytosis transcriptional programs in macrophage populations

We recently characterized cellular heterogeneity within the early wound beds of mice and found that fibroblasts, dendritic cells, and neutrophils all strongly expressed genes associated with cell death, apoptosis, and apoptotic recognition receptors, suggesting that these cells are undergoing apoptosis and performing efferocytosis in these early time points (Justynski et al., 2023). To further characterize the transcriptional response of efferocytic macrophages on macrophage phenotypes, we sought to determine how macrophage heterogeneity within skin wounds related to the in vivo and in vitro gene programs of macrophages performing efferocytosis.

Focusing on macrophage heterogeneity within scRNA sequencing data of mid-stage skin wounds (Haensel et al., 2020), we could broadly define 3 sets of macrophages including inflammatory macrophages that express *Ccl8, Ccl12, Mrc1,* and *Ccr2*, resolving macrophages with high *Vegfa*, *Fn1, Arg1,* and *Cd9* expression, and a mixed population that expressed inflammatory and resolving markers (**Figs. 4a and 4b**). To determine if the efferocytosis gene signatures mapped onto specific macrophages subsets, we calculated expression-based scores for the top 50 highly expressed and upregulated genes in Eff+ cells in vivo and applied them to the scRNA-seq data. The Eff+ signature was enriched in macrophages with the resolving and mixed transcriptional program with little enrichment in the inflammatory cluster. The expression of two highly upregulated genes in the Eff+ in vivo signature, *Srgn* and *Slpi*, highlights this preferential enrichment of Eff+ macrophage gene expression in the resolving macrophage subsets (**Fig. 4e**).

**Figure 4:**
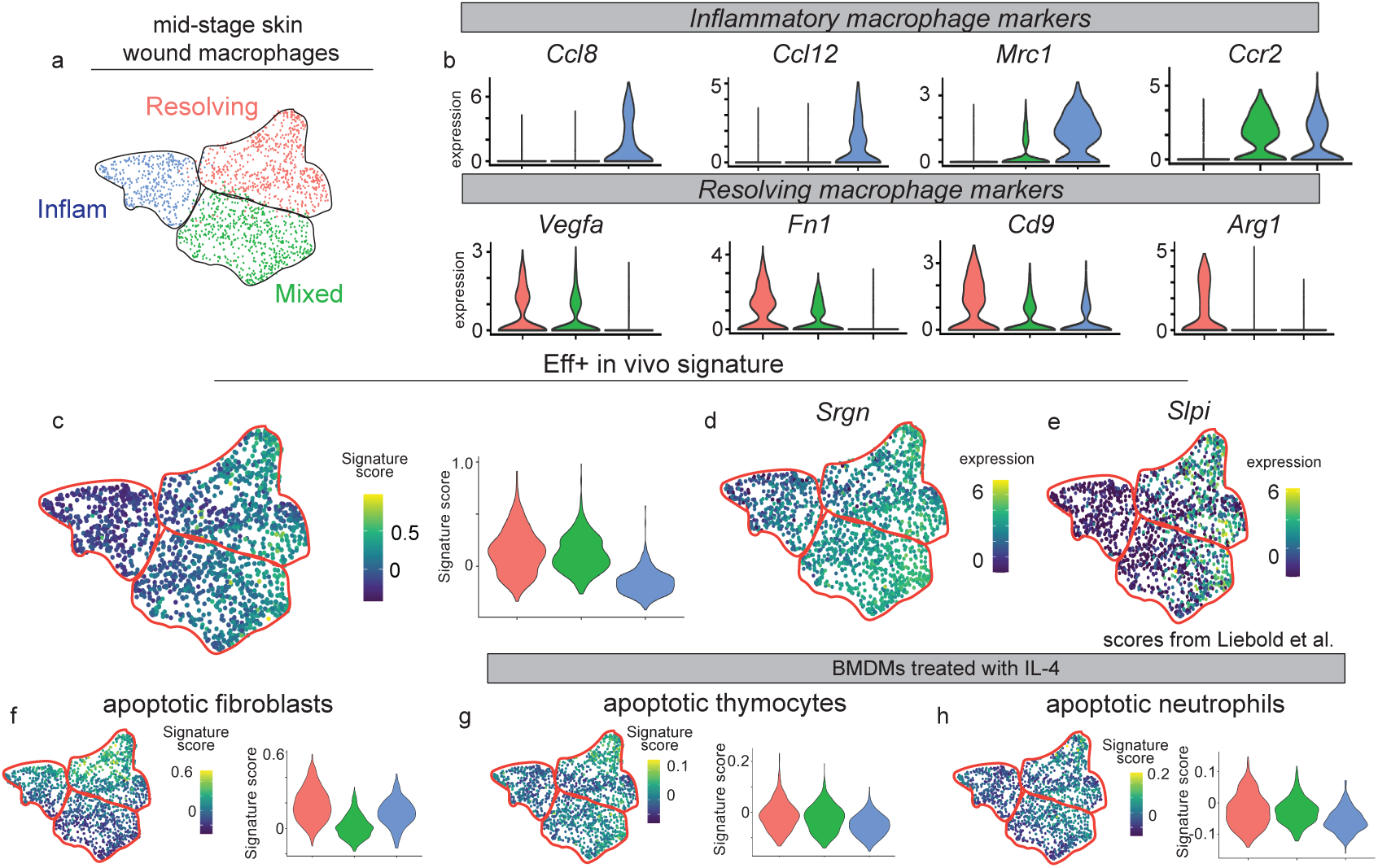
Macrophage efferocytosis gene signatures map onto heterogeneous macrophage populations within skin wounds. a) UMAP plot of single-cell RNA sequencing (scRNA-seq) data for macrophages from mid-stage wound beds in murine back skin. Data is from Haensel et al. 2020. b) Violin plots of differentially expressed marker genes of inflammatory and resolving macrophages in mid-stage wound beds. c) UMAP and violin plot of in vivo Eff+ gene signature score in macrophages from mid-stage wound beds in murine back skin. d-e) UMAP plots of *Srgn* and *Slpi* expression in macrophages from mid-stage wound beds in murine back skin. f-h) UMAP and violin plot of gene signature scores from apoptotic fibroblasts (f), apoptotic thymocytes (g) or neutrophils (h) from mid-stage wound beds in murine back skin.

To determine whether transcriptional changes in macrophages that efferocytosed apoptotic fibroblasts were similar to specific macrophage subsets, we calculated expression-based scores for the top 50 differentially upregulated and highly expressed genes from the Eff+ in vitro data (**Fig. 3**) and found that the apoptotic fibroblast signature mapped onto inflammatory and resolving macrophages with low expression in the mixed macrophage population (**Fig. 4c**). Using a Rank Rank Hypergeometric Overlap (RRHO) analysis, which evaluates similarities between gene expression profiles (Plaisier et al., 2010), we found that few shared genes exist between the Eff+ transcriptome from skin wounds and Eff+ gene expression from BMDMs treated with apoptotic fibroblasts (**Fig. S5**).

Recent work demonstrated that apoptotic neutrophils, hepatocytes, and T cells could impart differential gene expression changes on efferocytosing macrophages after IL-4 induction (Liebold et al., 2024), suggesting that apoptotic cell identity can influence macrophage phenotypes. To further analyze whether the apoptotic neutrophils or thymocytes shared a transcriptional signature with macrophage subsets in skin wounds, we mapped the gene signatures of BMDMs incubated with apoptotic neutrophils and thymocytes and stimulated with IL-4 (Liebold et al., 2024) onto the scRNA sequencing data from skin wound macrophage populations. Both the apoptotic thymocyte and neutrophil signatures had a low signature score but were more enriched in the resolving and mixed macrophage subsets. Taken together, these data indicate that macrophage efferocytosis induces a resolving transcriptional profile.

## Discussion

In this paper, we define how efferocytosis impacts macrophage transcription within mouse skin wounds. We found that efferocytic macrophages within mid-stage skin wounds upregulate genes associated with blood vessel repair, epithelial cell proliferation, and inflammatory signaling rather than traditional efferocytosis pathways that are upregulated in in vitro efferocytosis assays such as actin, chemotaxis, and phagocytosis. Consistent with the upregulation of genes involved in tissue repair, rather than inflammation, we find that gene signatures of macrophages undergoing efferocytosis map onto resolving macrophages within scRNA sequencing data of skin wounds. Thus, our data support the existing model that efferocytosis promotes tissue repair and resolution of inflammation and resonates with prior work demonstrating the importance of the tissue environment on macrophage heterogeneity induced by efferocytosis (Gonzalez et al., 2017).

Our study also supports previous research analyzing the transcriptional response induced by macrophage efferocytosis. We find that efferocytosis+ macrophages in vivo upregulated cell proliferation pathways, similar to recent work revealing the importance of efferocytosis on macrophage proliferation in an inflammatory environment (Ngai et al., 2023). We also found a broad expression regulation of metabolic changes, including nucleotide and phosphate metabolic processes, which have been implicated previously in the mechanism for anti-inflammatory transition due to efferocytosis (Schilperoort et al., 2023). In vitro, we found upregulation of Mfge8 and Anxa1, both previously associated with efferocytosis and revascularization during wound healing (Cabrera & Makino, 2022; Das et al., 2016; Huang et al., 2020; Khanna et al., 2010). Our data adds to the growing evidence that apoptosis recognition may influence specific aspects of tissue repair including angiogenesis (Jin et al., 2024), which we found was abrogated in murine skin wounds in which Axl and Timd4, two receptors that mediate efferocytosis, were inhibited (Justynski et al., 2023; Li et al., 2010; Zhu et al., 2019).

Similar to prior studies in vitro, BMDM efferocytosis of apoptotic fibroblasts led to an upregulation of genes associated with actin remodeling, phagocytosis, and chemotaxis (Larson et al., 2016; Mota et al., 2021). Interestingly, efferocytosis of apoptotic fibroblasts induced genes that mapped onto immature macrophages, which may indicate that these are early efferocytosis response genes. Consistent with this hypothesis, macrophages that efferocytose apoptotic neutrophils and were treated with IL-4, which promotes macrophage differentiation, upregulate tissue repair genes (Bosurgi et al., 2017), and are transcriptionally similar to resolving macrophages in skin wounds, rather than inflammatory macrophages. While our in vivo reporter line doesn’t distinguish between different apoptotic cellular targets that might alter gene expression in vivo (Liebold et al., 2024), mid-stage wounds harbor monocyte-derived macrophages/dendritic cells, fibroblasts, keratinocytes, and endothelial cells that might undergo apoptosis and could alter macrophage phenotypes (Haensel et al., 2020; Wasko et al., 2022). Future work defining how the identity of apoptotic skin cell types influences macrophage phenotype and function may reveal additional functions of efferocytosis in skin wounds.

Several other possible differences between the in vitro and in vivo models could explain the differences in gene expression. Beyond different cell identities, different types of cell death can also alter macrophage phenotypes (Naessens et al., 2024; Shiratori-Aso et al., 2023) and could change macrophage function in vivo. Neutrophil extracellular traps, ferroptosis, and pyroptosis have been shown to contribute to chronic diabetic wounds and can be present in non-diabetic wounds (Huang et al., 2023; Mu et al., 2022). These alternative types of cell death have been shown to produce different inflammatory effects in nearby phagocytosing macrophages (Naessens et al., 2024; Shiratori-Aso et al., 2023). Second, the in vivo wound niche likely contains signals for macrophage differentiation which are absent in our in vitro experiments and can alter macrophage phenotype (Aitcheson et al., 2021). A caveat of our in vivo data set is that only uptake of non-myeloid cells that are TdTom+ are detected, while apoptosis of myeloid lineage cells such as macrophages and dendritic cells, as well as neutrophils that express LysM are likely a large proportion of apoptotic cells within skin wounds and the impact of their efferocytosis would be omitted from our data sets.

In summary, our data indicate that efferocytosis drives macrophages towards a resolving phenotype and suggests that efferocytosis may directly regulate angiogenesis and keratinocyte proliferation to drive tissue repair. Interestingly, transplantation of blood-derived macrophages into skin wounds can promote healing (Danon et al., 1997), and this effect may improve with efferocytosis stimuli (Liebold et al., 2024). Given the importance of apoptosis in wound repair and defects in apoptosis recognition receptor signaling within human diabetic wounds, future work exploring how macrophage efferocytosis controls wound repair may reveal therapeutic tissue repair mechanisms for chronic skin wounds.

## Methods

### LysMCreER/mTmG mouse

All experimental procedures were approved and in accordance with the Institutional Animal Care and Use Committee.WT C57BL/6J mice (Strain #:000664), Lyz2tm1(cre/ERT2)Grtn/J (LysMCreER) mice (Strain #:031674), and B6.129(Cg)-Gt(ROSA)26Sortm4(ACTB-tdTomato,-EGFP)Luo/J (mTmG) mice (Strain #:007676) were purchased from The Jackson Laboratories. LysMCreER/mTmG mouse lines were generated by our lab from these purchased lines. Mice were housed in an Association for Assessment and Accreditation of Laboratory Animal Care (AALAC)-accredited animal facility at Yale University (Protocol # 11248). Animals were maintained on a standard chow diet ad libitum (Harlan Laboratories, 2018S) in 12 hr light/dark cycling. Up to five mice were housed per cage. For experiments using intraperitoneal (i.p.) tamoxifen administration, 100 μL of 30 mg/mL tamoxifen (Sigma-Aldrich) in sesame oil was injected daily for 3 days prior to experiments (Shook et al., 2020).

### Wounding

Seven- to 9-week-old male mice were wounded during the telogen phase of hair cycling, which was confirmed by the pink skin color of pigmented mice. Mice were anesthetized using isoflurane and six full-thickness wounds, at least 4 mm apart, were made on shaved back skin using a 4 mm biopsy punch (Millitex), as described previously (Shook et al., 2020). Animals were sacrificed at noted intervals after injury and wound beds were processed for subsequent analysis.

### Flow cytometry/FACS

Wound beds were digested for further flow cytometry analysis or sorting in a buffer of Roswell Park Memorial Institute (RPMI) medium with glutamine (Gibco), Liberase Thermolysin Medium (TM) (Roche), DNase, N-2-hydroxyethylpiperazine-N-2-ethane sulfonic acid (Gibco), sodium pyruvate (Gibco), non-essential amino acids (Gibco), and antibiotic-antimycotic (100X) (Gibco). Digested wounds or cultured cells were blocked with 3% Bovine Serum Albumin in PBS. Cells were stained with antibodies for 30 min on ice. Macrophages were defined as F4/80+/CD45+ cells. In vivo efferocytosis+ macrophages from LysMCreER/mTmG mice were defined as F480+/CD45+/TdTomato+/GFP+ cells, and efferocytosis− macrophages were defined as F480+/CD45+/TdTomato-/GFP+ cells. In vitro experiments using primary skin fibroblasts defined efferocytosis+ macrophages as F4/80+/CD45+/TdTomato+, while efferotycosis− macrophages were F480+/CD45+/TdTomato-. All sorting was performed with a FACS Aria III with FACS DiVA software (BD Biosciences). For Amnis flow cytometry, digested wounds were fixed with 4% PFA for 15 min prior to antibody incubation. AMNIS experiments were performed at the Yale Flow Cytometry Center using a Cytek^®^ Amnis^®^ ImageStream^®X^ MKII. Flow cytometry analysis was performed using FlowJo Software (FlowJo) for FCS files and IDEAS software for AMNIS analysis. Antibodies used are shown in Table S7.

### Quantitative real-time PCR

Cells isolated by FACS were digested using TRIzol LS (Invitrogen). RNA was extracted using the RNeasy Plus Mini Kit (QIAGEN) per manufacturer’s instructions. cDNA was generated using equal amounts of total RNA with the Superscript III First Strand Synthesis Kit (Invitrogen) per manufacturer’s instructions. All quantitative real-time PCR was performed using SYBR green on a LightCycler 480 (Roche). Primers for specific genes are listed in Table S7. Results were normalized to β-actin.

### RNA seq analysis

RNA was isolated from FACS sorted macrophages using the RNeasy Plus Mini Kit (QIAGEN). RNA sequencing was performed by the Yale Center for Genome Analysis. With trimmomatic version=0.39, Clontech adapters were trimmed, leading and trailing low quality bases were removed, and reads below the 36 bases long were dropped. Reads were aligned to genome Mus musculus. GRCm39 with Star RNA-aligner version=2.7.10a. With the Star option --quantMode GeneCounts, the number of reads per gene were counted. All gene counts with a sum (over all samples) of less than 500 were filtered out. Then the differential expression analysis was performed with DESeq2. Gene ontology analysis was performed using Panther (Thomas et al., 2022).

For analysis of GSE142471 single cell RNA sequencing data (Haensel et al., 2020), we used Seurat (v5.1) (Stuart et al., 2019). After normalization, finding variable features, and scaling the data, we used UMAP dimension reduction for visualization and Louvain clustering based on their first 10 principal components. Louvain clustering was run at a resolution of 0.2. For marker detection, we used the Wilcoxon rank sum test, comparing one cluster’s cells against all remaining cells. For further analysis, we repeated the previous steps, starting with the normalization on the macrophage cluster, which expressed high levels of *Adgre1* (F4/80) and *Cd68*. We used the *AddModuleScore* function to calculate four gene expression scores based on the previously described Eff+ in vivo, aF, IL-4–aH, IL-4–aT treatment-specific gene lists for each cell (Table S8).

### Rank-Rank Hypergeometric Overlap

The extent of overlap between in vivo and in vitro RNA sequencing sets was compared using Rank-Rank Hypergeometric Overlap (RRHO) (v. 1.44.0) as described previously (Cahill et al., 2018). To perform RRHO analysis, the RNA seq lists were first ranked by −log10(pvalue) according to differential seq analysis. The RRHO algorithm was then used to find overlap in thresholds of ranked differential expression between different studies.

### BMDM differentiation and isolation

Femurs and tibia from WT C57BL/6J mice were dissected and cleaned. Bones were then cut open, and 1mL of RPMI with 15% Newborn Calf Serum (NCS) was flushed through the bone over a 40 uM mesh to remove marrow. After centrifugation, cell pellets were treated with ACK lysis buffer for 1 min. Bone marrow cells were counted, and cultured in a media containing 15% NCS, 1% antifungal antibiotic, and 10ng/mL GMCSF for one week. Resulting differentiated macrophages were used for further experiments.

### Fluorescence imaging

Mouse wound beds were embedded in optimum cutting temperature compound (VWR) and wound beds were sectioned through their entirety to identify the center. 12µm cryosections were fixed with 4% formaldehyde and immunostained as previously described with the following antibodies: GFP (chicken, 1:1,000; Abcam) and RFP (Rabbit, 1:400, Rockland). Composite images were acquired using the tiles module on a Zeiss AxioImager M1 (Zeiss) equipped with an Orca camera (Hamamatsu).

### Primary Fibroblast isolation and culture

Skin of mTmG+ mice was digested in a buffer for isolation with DMEM High Glucose (Gibco), Liberase Thermolysin Medium (TM) (Roche), DNase, N-2-hydroxyethylpiperazine-N-2-ethane sulfonic acid (Gibco), sodium pyruvate (Gibco), non-essential amino acids (Gibco), and antibiotic-antimycotic (100X) (Gibco). Digested tissue was filtered through 100µm meshes, and cell suspensions were cultured in DMEM F-12 (ATCC) with 10% fetal bovine serum (FBS) and 1% antifungal antibiotic and passaged at least twice to ensure only proliferative fibroblasts remained. Apoptosis was induced by 30 min UV exposure, followed by 24 hrs of incubation. Cell death was validated by assaying % live cells using a Countess 3 (Thermo Fisher).

### Statistics

To determine significance between two groups, comparisons were made using Student’s t-test. Analyses across multiple groups were made using a one- or two-way ANOVA with Bonferroni’s post hoc using GraphPad Prism for Mac (GraphPad Software, La Jolla, CA, USA) with significance set at p<0.05. Sample sizes were determined using power analysis and taking into consideration our experience with the wounding model.

## Data Availability Statement

Sequencing and analysis data can be found in NCBI’s Gene Expression Omnibus (GEO) under accession number <u>GSE273754 (</u>Bulk RNA seq of efferocytosis in vivo and in vitro) and <u>GSE142471</u> (mid-stage wound scRNA sequencing data) (Haensel et al., 2020). All other data is available upon request.

## Author Contributions Statement

Conceptualization: WK, VH

Formal Analysis: WK, DK, VH

Funding acquisition: VH

Investigation: WK

Software: DK

Visualization: WK, VH

Writing – Original Draft Preparation: WK, VH

Writing – Review and Editing: WK, VH

## Conflict of Interest

The authors declare no conflicts of interest.

## Declaration of Artificial Intelligence/Large Language Model use

During the preparation of this work the authors used ChatGPT for assistance with basic R code and commands during data analysis. After using this tool/service, the authors reviewed and edited the content as needed and take full responsibility for the content of the publication.

## Acknowledgements

We would like to thank members of the Horsley laboratory for their feedback and critical analysis of these data and manuscript. We would also like to thank the Yale Animal Resources Center (YARC) staff for animal husbandry, Yale School of Medicine and Yale Science Hill Flow Cytometry Core, and Yale Center for Genome Analysis, which receives funding from NIH Shared Equipment grant #1S10OD028669-01. W.K. received funding from the Rick Li and Yanning Sun Fellowship Fund. V.H. is funded by N.I.H-NIAMS R01s AR076938, AR0695505, and AR075412.

---

**Supplementary figure 1:**
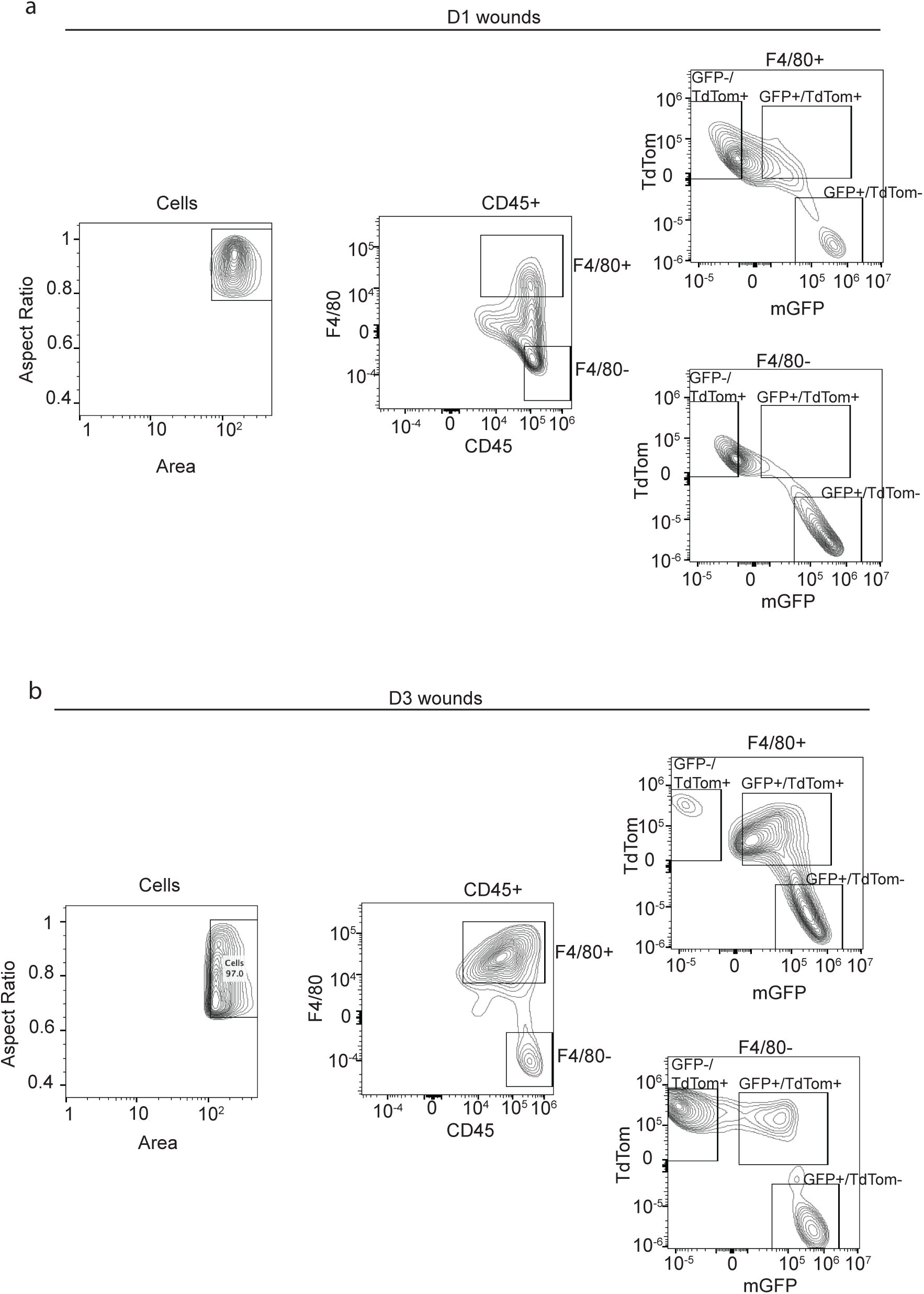
AMNIS flow cytometry gating strategy of D1 and D3 wounds. a) Gating strategy for AMNIS flow cytometry analysis of D1 wounds. Gates were Area/Aspect ratio to determine cells, CD45/F4/80 to determine immune cells and macrophages, TdTom/GFP from the LysMCreER/mTmG mouse line. b) Gating strategy for AMNIS flow cytometry analysis of D3 wounds. Gates used were Area/Aspect ratio to determine cells, CD45/F4/80 to determine immune cells and macrophages, TdTom/GFP from the LysMCreER/mTmG mouse line.

**Supplementary figure 2:**
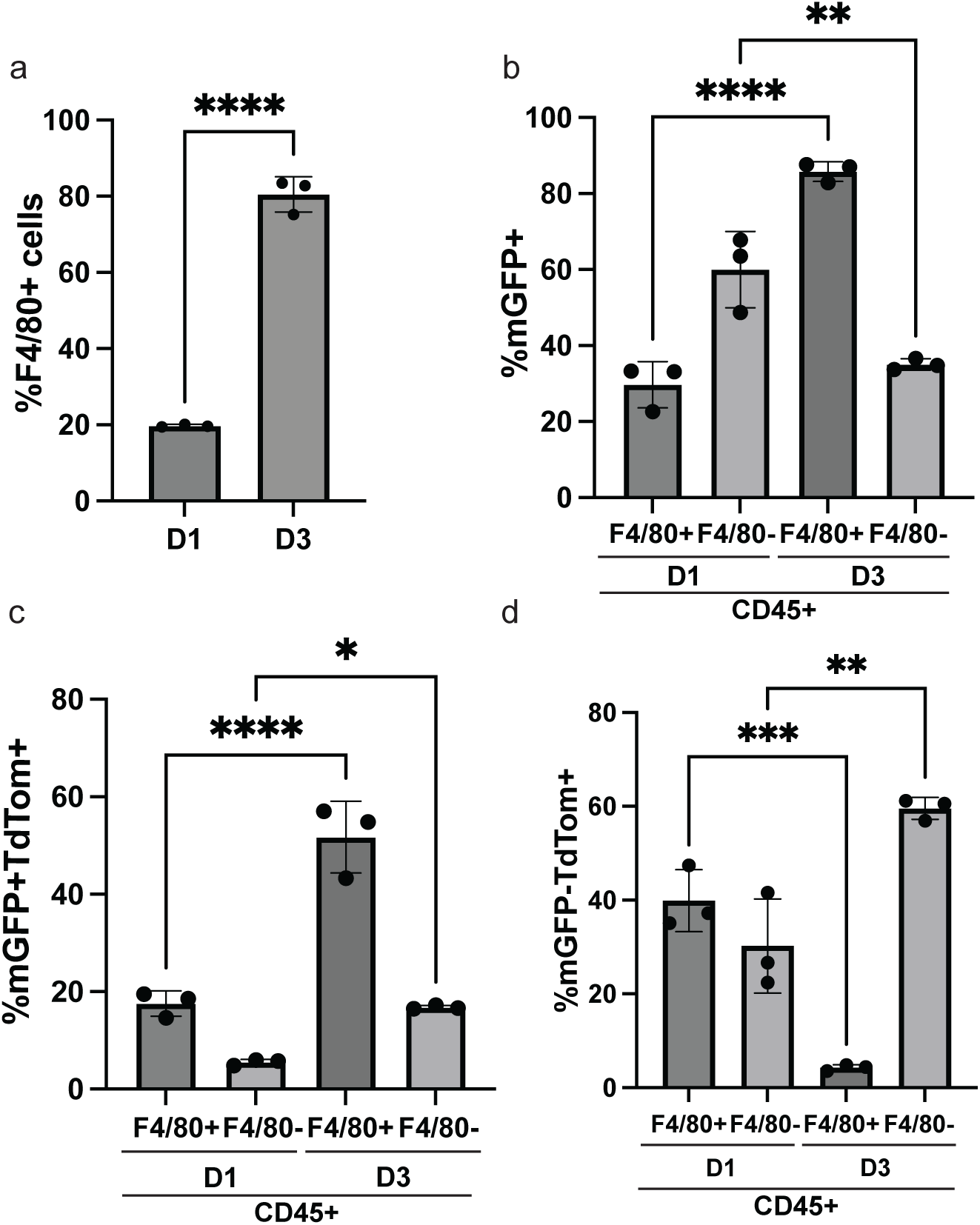
Quantification of Flow Cytometry analysis from D1 and D3 wounds. a) Quantification of % F4/80+ cells via AMNIS flow cytometry from D1 and D3 mouse wounds. N=3 mice. b) Quantification of % GFP+ cells in CD45+ populations from D1 and D3 mice. N=3 mice. c) Quantification of % GFP+TdTom+ cells in either CD45+/F4/80+ or CD45+/F4/80− populations as labeled in D1 and D3 mice. N=3 mice. d) Quantification of % GFP-TdTom+ cells in either CD45+/F4/80+ or CD45+/F4/80− populations as labeled in D1 and D3 mice. N=3 mice. Error bars indicate mean +/− SEM. p values calculated with one way ANOVA. *p<.05, **p<.005, ***p<.0005, ****p<.00005

**Supplementary figure 3:**
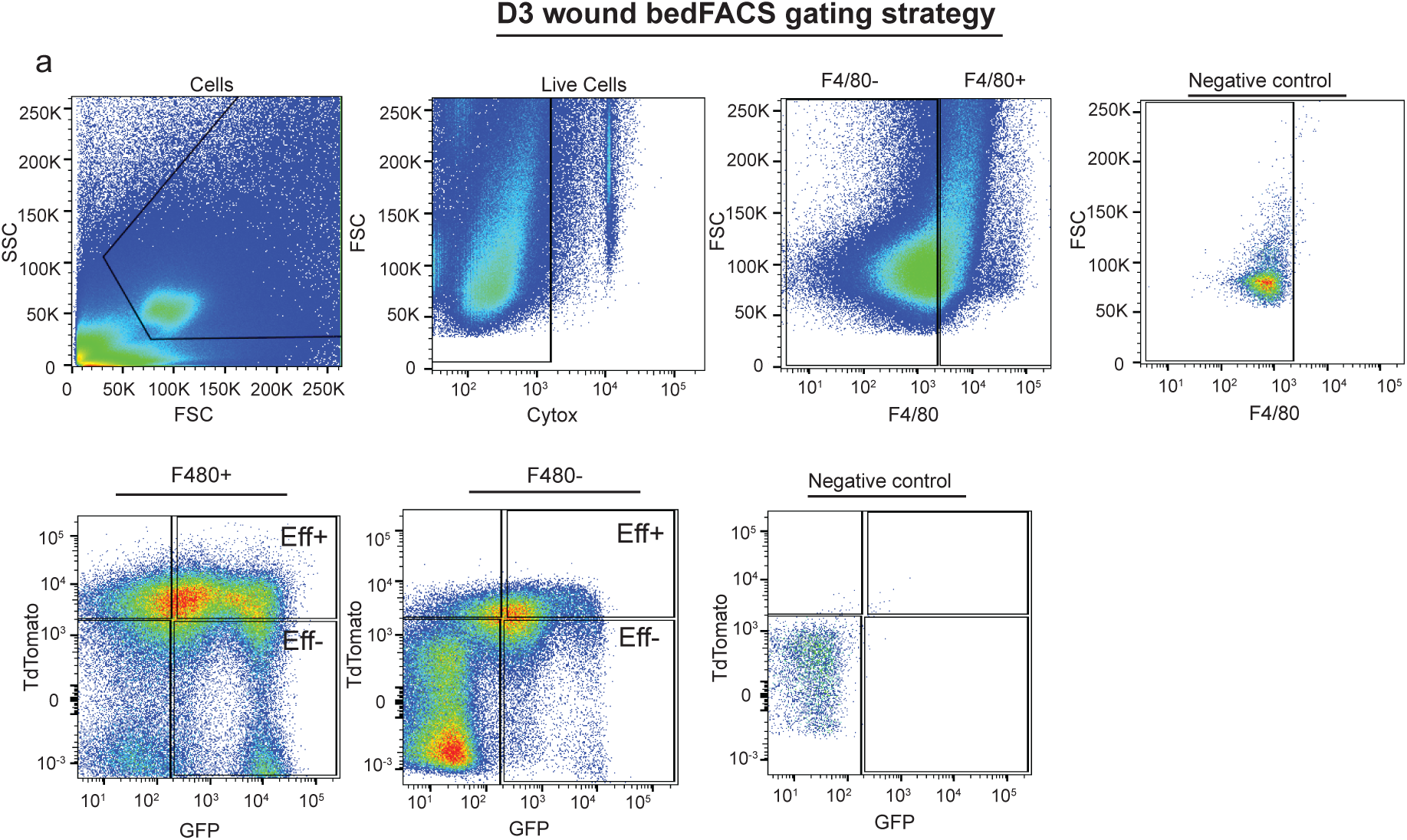
Gating strategy for ARIA fluorescence associated cell sorting from D3 wounds. a) Gating strategy for ARIA flow cytometry sorting of macrophages from D3 wounds. First cells were sorted via FSC and SSC, then live cells as Cytox-, then macrophages as F4/80+, then Eff+ as TdTom+GFP+ and Eff− as TdTom-GFP+. As labeled, F4/80− sorted cells are also shown for GFP and TdTom analysis. Unlabeled cells derived from mouse spleens are shown as negative control.

**Supplementary figure 4:**
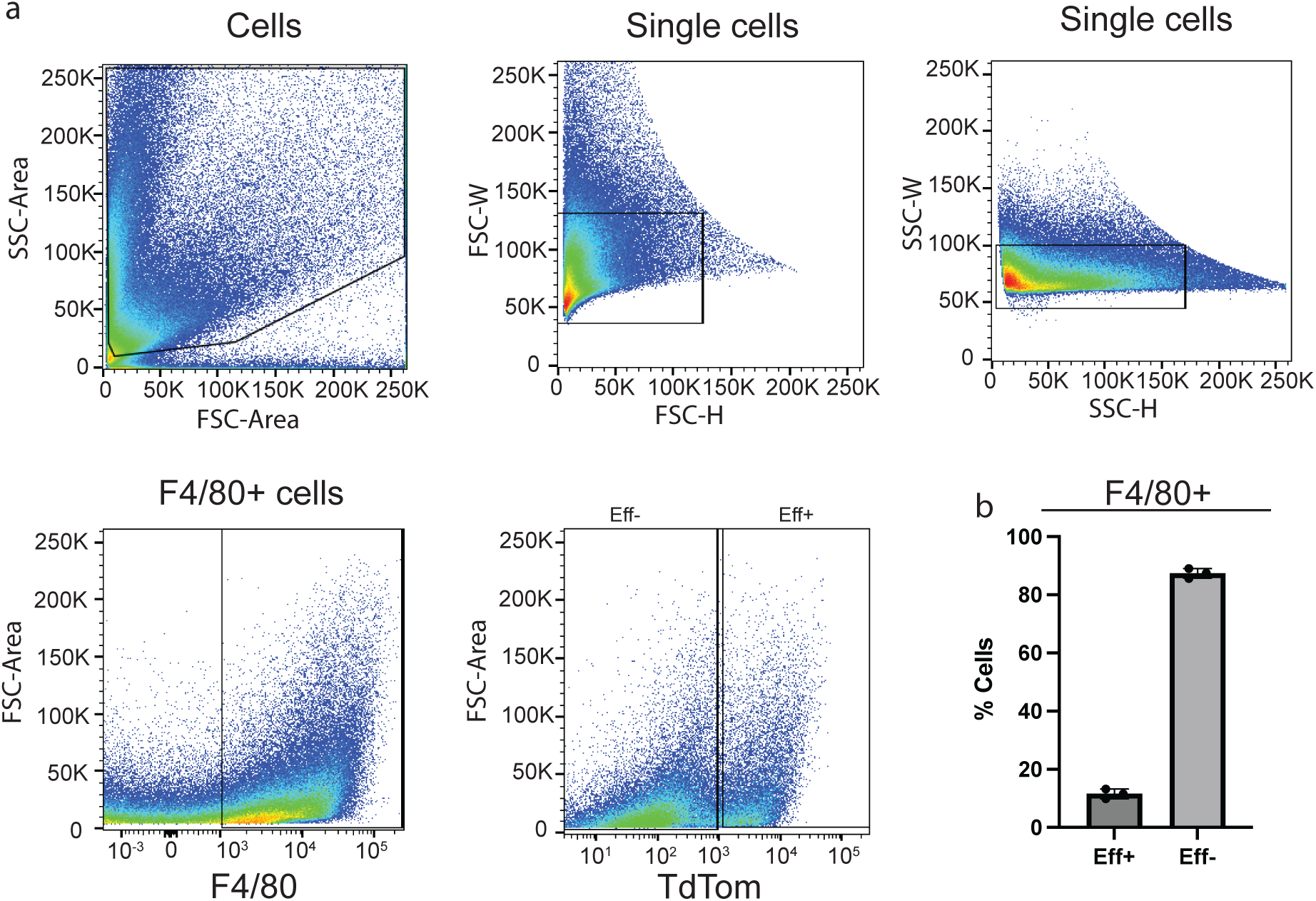
Gating strategy for ARIA fluorescence associated cell sorting from in vitro efferocytosis. a) Gating strategy for ARIA flow cytometry sorting of in vitro macrophages performing efferocytosis. Cells were sorted from FSC and SSC, single cells were purified by FSC-H/FSC-W and SSC-H/SSC-W, macrophages were sorted by F4/80+, and Efferocytosis+ cells were determined as TdTom+. b) Quantification of Eff+ cells. Error bars indicate mean +/− SEM. p values calculated with one way ANOVA.

**Supplementary figure 5:**
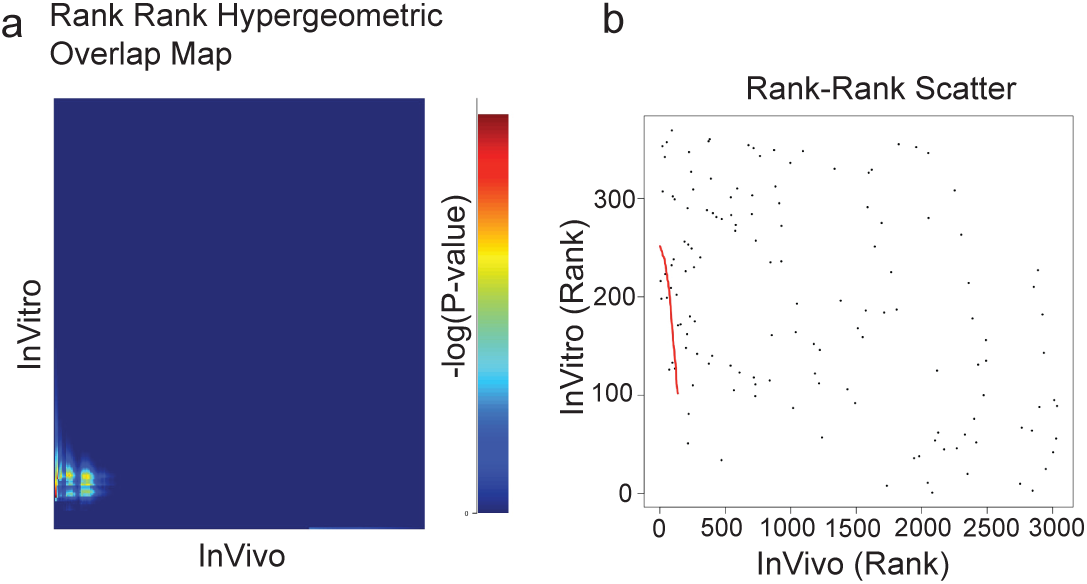
RRHO comparison between in vivo and in vitro efferocytosis. a) Rank Rank Hypergeometric Overlap (RRHO) test of overlap between in vivo and in vitro RNA-seq data for all differentially genes. b) Scatterplot showing rank-rank comparisons of all genes between in vivo and in vitro RNA-seq data. Red line indicates line of best fit.

**Supplementary table 1: Differentially regulated genes identified between in vivo Eff+ and Eff− cells**

**Supplementary table 2: Differentially regulated genes identified between in vitro Eff+ and Eff− cells**

**Supplementary table 3: Gene ontology analysis of genes upregulated in Eff+ cells in vivo**

**Supplementary table 4: Gene ontology analysis of genes downregulated in Eff+ cells in vivo**

**Supplementary table 5: Gene ontology analysis of genes upregulated in Eff+ cells in vitro**

**Supplementary table 6: Gene ontology analysis of genes downregulated in Eff+ cells in vitro**

**Supplementary table 7: List of primers and antibodies used**

**Supplementary table 8: List of genes used to determine gene signature for Figure 4**

## Notes

https://www.ncbi.nlm.nih.gov/geo/query/acc.cgi?acc=GSE273754

https://www.ncbi.nlm.nih.gov/geo/query/acc.cgi?acc=GSE142471

### Competing Interest Statement

The authors have declared no competing interest.

